# Ribosome biogenesis is a therapeutic vulnerability in paediatric neuroblastoma

**DOI:** 10.1101/2025.03.26.645392

**Authors:** Camille Jouines, Piero Lo Monaco, Angéline Gaucherot, Marie-Ambre Monnet, Isabelle D. Iacono, Valentin Simioni, Déborah Monchiet, Jean-Jacques Diaz, Valérie Combaret, Virginie Marcel, Frédéric Catez

**Author notes:** These authors equally contributed to the study (Virginie Marcel, Frédéric Catez).

## Abstract

**Background:** Neuroblastoma is a heterogeneous malignant paediatric tumor with prognosis depending on patient age and disease stage. Current treatment strategies rely on four key diagnostic criteria: age, histological stage, *MYCN* gene status, and genomic profile. It has been reported that MYC oncogenic activity depends on ribosome biogenesis, whose hyperactivation in cancer cells supports their high proliferative capacity, and thus represent a potential therapeutic target.

**Methods:** we utilized the well-established IMR-32 cell line along with a panel of patient-derived neuroblastoma cell lines with varying MYCN status, which we previously established. Additionally, we generated an IMR-32 cell line expressing an shRNA targeting the ribosome biogenesis factor fibrillarin (FBL). Cell growth, apoptosis markers, and cell cycle regulators were analyzed. Expression of ribosome biogenesis factors was assessed using publicly available datasets and RT-qPCR data from an in-house neuroblastoma cohort.

**Results:** We explored whether ribosome biogenesis represents a vulnerability in neuroblastoma. Our findings demonstrate that inhibition of RNA polymerase I using CX-5461 and BMH-21 suppressed cell proliferation at nanomolar concentrations and induced ribosomal stress, leading to activation of apoptosis and the p21 pathway. Furthermore, we identified FBL as a marker of poor prognosis in neuroblastoma. Consistently, FBL knockdown reduced neuroblastoma cell proliferation, supporting its potential as a therapeutic target.

**Conclusion:** Our study reinforces the therapeutic potential of ribosome biogenesis inhibition in neuroblastoma and expands the list of potential targets to include rRNA maturation factors. These findings highlight the promise of targeting ribosome biogenesis as a novel approach for neuroblastoma treatment.

## Introduction

Neuroblastoma is a malignant paediatric tumor of the peripheral sympathetic nervous system, originating from immature foetal nerve cells. Neuroblastoma is the most common extracranial solid tumor in children, making it the third most prevalent paediatric cancer [1], with 90% of cases diagnosed before the age of 5-years-old [2]. Neuroblastoma accounts for 7% of malignancies in patients under 15-years-old and approximately 15% of all cancer-related deaths in paediatric populations [3]. Despite recent therapeutic advancements, 5-year survival rates for high-risk pediatric neuroblastoma has remained at approximately 50% over the past two decades [4], emphasizing the need for continued research on this paediatric cancer.

Neuroblastoma is a clinically heterogeneous disease requiring treatments adapted to individual risk. Indeed, prognosis mainly relies on patient age and disease stage at diagnosis [1], [5]. Poorest survival is mainly observed for children diagnosed with advanced stage 4 disease at > 18 months. In 2015, Pinto *et al*. described a classification system termed the International Neuroblastoma Risk Group (INRG), which uses combinations of prognostic risk factors to define distinct risk groups. Based on 5-year event-free survival (EFS) rates, patients are categorized into very low, low, intermediate, or high-risk groups. The INRG classification considers imaging criteria, the extent of locoregional disease, and the presence of disseminated metastases (skin, liver, and bone marrow) [5]. In the low or intermediate-risk groups, neuroblastoma patients achieve excellent outcomes, with long-term survival rates exceeding 90% [6]. In low-risk cases, surgery alone, even if incomplete, has proven curative for nearly all patients. In certain subgroups, spontaneous tumor regression allows for a cure without any treatment [6]. For high-risk patients, current treatment involves intensive multimodal therapy: five to six cycles of induction chemotherapy and surgery, consolidation therapy with high-dose of chemotherapy and radiation, and post-consolidation therapy targeting residual cells. These high-risk neuroblastoma patients have a 5-year overall survival of only 40% to 50% [6], [3]. Therefore, new therapeutic advances, such as CAR T-cell therapy, are also being explored to fill the urgent need for novel treatments in these challenging paediatric cancer types [5]. In parallel, early detection of the most aggressive neuroblastoma tumours, is also required to improve outcomes.

In addition to the INGR classification, genetic features such as ploidy, *MYCN* copy number, and 1p or 11q deletions allows distinguishing aggressive tumors from those likely to regress. For example, the identification of *MYCN* amplification, observed in approximately 25% of cases [2], serves as an independent predictive marker of poor prognosis, associated with invasive and metastatic behavior. Thus, in addition to age and histological stage, *MYCN* gene status and genomic profile are now evaluated at diagnosis to assess cure prospects and guide therapeutic decision [3]. More importantly, these discoveries have led to novel strategies for neuroblastoma treatment. For instance, in cases of relapse, the *ALK* gene status can help assess the interest of targeted therapy. Unfortunately, MYC targeting remains challenging, necessitating the exploration of alternative approaches, such as targeting MYC-driven pathways [5], [2].

MYC is major activator of ribosome biogenesis, by increasing activity of the RNA polymerase I (RNA Pol I), and regulating expression of a large set of ribosomal proteins and ribosome biogenesis factors. Consequently, MYC protein overexpression increases ribosomal RNA (rRNA) synthesis [7]. Consistently, in neuroblastoma, tumours with unfavorable clinical outcome are characterized by overexpression of genes driving ribosome biogenesis and protein synthesis, supporting the notion than *MYCN*-driven neuroblastoma might particularly rely on ribosome biogenesis [1]. More importantly, it has been shown that MYC oncogenic activity relies on ribosome biogenesis, and that impeding MYC-driven ribosome biogenesis overactivation is sufficient to reduce MYC-driven leukemogenesis [8]. Thus, inhibiting ribosome biogenesis appears as a novel approach to treat MYC-driven tumors.

First evaluations of ribosome biogenesis inhibition have been conducted in neuroblastoma using RNA Pol I inhibitors including Actinomycin D, and CX-5461 and CX-3543. Actinomycin D (ActD) and CX-5461 have been shown to induce cell death in vitro and reduce tumor progression in vivo [9], [10], [4]. Yet, the contribution of MYCN status to neuroblastoma cells response to ribosome biogenesis remains unclear, as studies reported conflicting results [6], [4]. Thus, RNA Pol I inhibitors are promising targeted therapies in neuroblastoma, including MYC-driven tumors. The two most characterized inhibitors, Actinomycin D and CX-5461, were shown to induce DNA damage, an activity that induces non-selective toxicity, and which require the development of new Pol I inhibitors or identification of novel targets in the ribosome biogenesis pathway.

Here, we evaluate BMH-21, another RNA Pol I inhibitor, for the first time in neuroblastoma using patient-derived cell lines. In addition, we investigated FBL as a novel target to inhibit ribosome biogenesis.

## Results

### Neuroblastoma cell lines are sensitive to RNA Polymerase I inhibitors

The effect of RNA Pol I inhibitors on neuroblastoma was first evaluated in the commonly used *MYCN*-amplified neuroblastoma cell line IMR-32, (Table S1, Figure 1A). IMR-32 cells were treated for 96 h with RNA Pol I inhibitors, CX-5461 or BMH-21, using concentration range from 1 to 1000 nM, and cell proliferation was assessed by MTS assay. The IC50 values for CX- 5461 and BMH-21 were 173.6 nM (SD ± 61) and 202.3 nM (SD ± 30.6), respectively. Thus, IMR-32 cell viability is impaired at nanomolar concentration, similarly to what previously observed for CX-5461 [10]. No significant difference was observed between CX-5461 and BMH-21 in this cell line, indicating that the two RNA Pol I inhibitors display the same activity in IMR-32.

**Figure 1.**
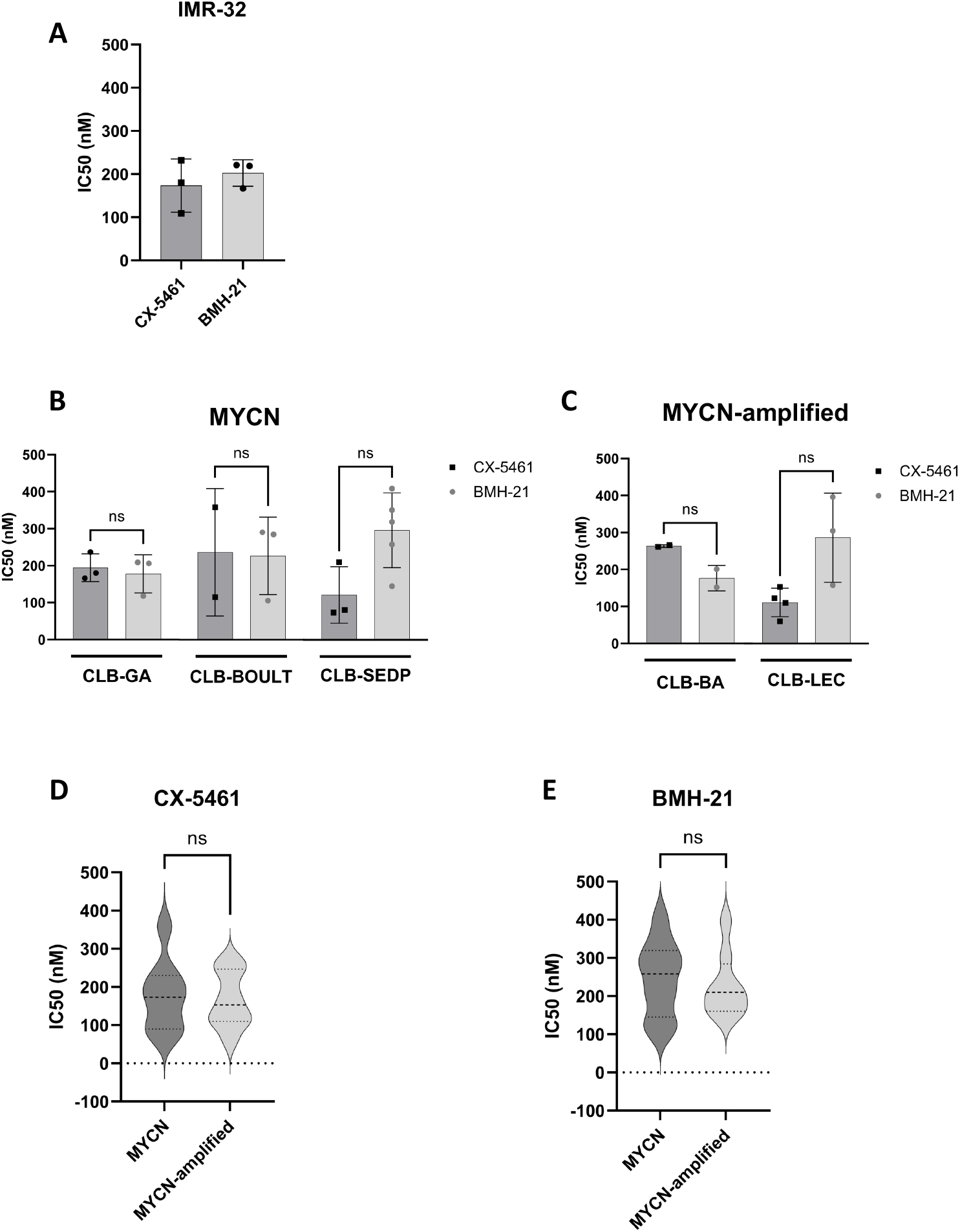
Sensitivity of neuroblastoma cell lines to RNA Pol I inhibitors. **(A-C)** IC50 values for CX-5461 and BMH-21 were determined by MTS assay at 96 h post-treatment with a 1 nM to 1µM concentration range, on IMR32 cell line (A) and five neuroblastoma patient-derived cell lines, with normal (B, D) or amplified (C, E) *MYCN* status. **(D-E)** The data of neuroblastoma patient-derived cell lines were pooled according to *MYCN* amplification status for CX-5461 (D) and BMH-21 (E) treatment. Each experimental condition was performed in at least two biological replicates. When more than 2 biological replicates were available, mean comparison was performed using Mann-Whitney test. ns = non significant.

We then compared the effect of these two RNA Pol I inhibitors in more relevant cancer models, using a panel of five neuroblastoma patient-derived cell lines [11]. Among these neuroblastoma patient-derived cell lines, two are *MYCN* -amplified (CLB-BA and CLB-LEC) (Table S1). Comparison of RNA Pol I inhibitors response using MTS assay 96 h post-treatment showed that IC50 values varied across different neuroblastoma patient-derived cell lines for both CX-5461 and BMH-21 (Figure 1B-C). For each neuroblastoma patient-derived cell lines, no significant difference in sensitivity was observed between CX-5461 and BMH-21 (Figure 1B-C), as observed for IMR-32. Notably, IC50 values were similar for both RNA Pol I inhibitors across the neuroblastoma patient-derived cell lines (Figure 1D-E), indicating that in this panel of neuroblastoma patient-derived cell lines, *MYCN* status did not modulate cell sensitivity to ribosome inhibition, in contrast to previous findings on CX-5461 in neuroblastoma [10]. Our data support previous observations that neuroblastoma cell lines, including patient-derived cell lines, are sensitive to the CX-5461 RNA Pol I inhibitor at nanomolar concentrations. Moreover, patient-derived neuroblastoma cell lines display similar sensitivity to the RNA Pol I inhibitor, BMH-21, independently of *MYCN* status.

### RNA Polymerase I inhibitors induces cell cycle arrest and apoptosis

In cancer cells, inhibition of ribosome biogenesis has been shown to promote a stress response termed ribosomal stress [12]. This stress response triggers cell cycle arrest and/or cell death. We recently reported that CX-5461 and BMH-21 promote cell cycle arrest but no cell death in triple-negative breast cancer [13]. To determine whether RNA Pol I inhibitors promote similar cellular ribosomal stress in neuroblastoma, we evaluated cell cycle arrest (cytostatic effect) and apoptosis (cytotoxic effect).

First, we analysed cell cycle inhibition by quantifying the mRNA levels of p21, a key cell cycle regulator and p53 effector, by RT-qPCR (Figure 2). IMR-32 and our panel of five neuroblastoma patient-derived cell lines were treated with 250 and 750 nM of CX-5461 and BMH-21 for 96 h. Actinomycin D (ActD) was used as a reference. In IMR-32, CX-5461 and BMH-21 treatment induced a 40 to 258-fold increase in p21 mRNA levels (Figure 2A). In our neuroblastoma patient-derived cell lines panel, a 4 to 87-fold in p21 mRNA level was observed (Figure 2B-2C). Interestingly, a dose-dependent induction of p21 mRNA levels was observed only in response to BMH-21 treatment. In addition, BMH-21 at 750 nM induced higher p21 mRNA levels than CX-5461 at 750 nM. Notably, similar increase in p21 mRNA levels was observed in non-amplified *MYCN* and *MYCN*-amplified cell lines (Figure 2B vs 2A-2C). These results suggest that in neuroblastoma cell lines, cell cycle arrest occurs in response to the two RNA Pol I inhibitors independently of *MYCN* status, BMH-21 triggering the strongest response.

**Figure 2.**
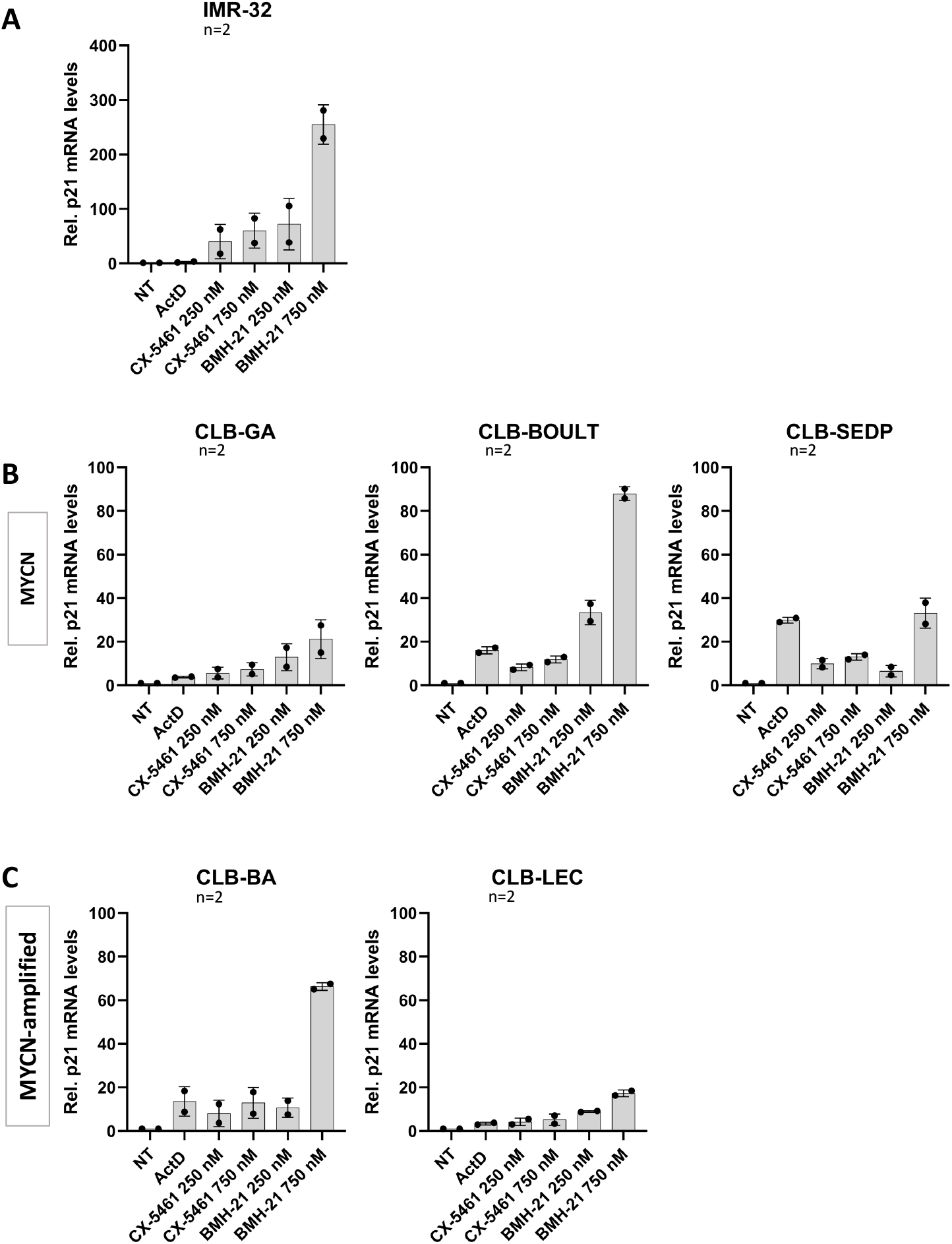
p21 mRNA levels in response to RNA Pol I inhibitors. p21 mRNA levels were quantified by RT-qPCR on cells treated with CX-5461 and BMH-21 (final concentrations: 250 nM and 750 nM) for 96 h in IMR32 cell line (A) and in neuroblastoma patient-derived cell lines, with either no *MYCN*-amplification (B) or *MYCN*-amplification (C). Actinomycin D (ActD) was used at 0.05 µg/mL. Data are mean values from 2 independent biological replicates.

Next, apoptosis was evaluated using a caspase 3/7 activity assay in our panel of neuroblastoma patient-derived cell lines in response to 96 h of treatment with CX-5461 and BMH-21 (Figure 3). In response to CX-5461 and BMH-21 treatment, a significant increase in caspase 3/7 activity was observed in IMR-32 and the 5 neuroblastoma patient-derived cell lines, although this activation was limited in the CLB-BOULT cell line (Figure 3B). In addition, CX-5461 promoted an higher induction of caspase 3/7 activity than BMH-21 in IMR-32, CLB-SEDP, CLB-BA and CLB-LEC lines, suggesting no correlation of CX-5421-induced caspase 3/7 activation and *MYCN* amplification status. Thus, in neuroblastoma cell lines, the apoptotic pathway is induced by both RNA Pol I inhibitors independently of *MYCN* status.

**Figure 3.**
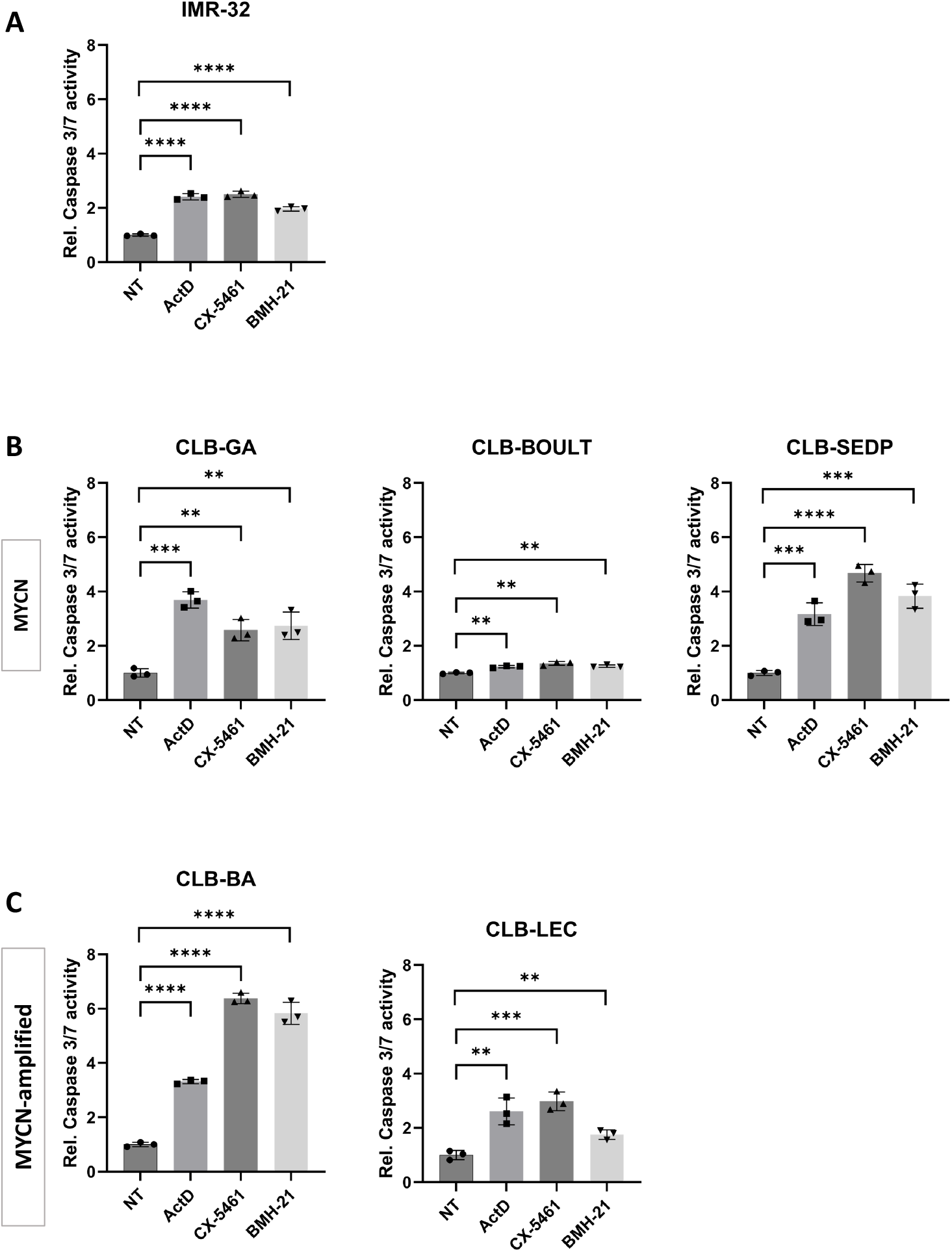
Caspase 3/7 activation in response to RNA Pol I inhibitors. Caspase 3/7 activation was assessed in response to CX-5461 and BMH-21 treatment at 500 nM) in IMR32 cell line (A) and in neuroblastoma patient-derived cell lines, with either no MYC-amplification (B) or MYC-amplification (C). Results are expressed as ratios relative to untreated cells. ActD: actinomycin D; Data are mean values from 2 independent replicates. ** p ≤ 0.01 ; *** p ≤ 0.001 ; **** p ≤ 0.0001

These findings demonstrate that RNA Pol I inhibitors promote both cell cycle arrest and cell death in neuroblastoma independently of *MYCN* amplification. Overall, these data support the efficacy of RNA Pol I inhibitors in neuroblastoma patient-derived cell lines.

### rRNA maturation factors are prognostic markers in neuroblastoma

In breast cancer, we recently made the proof-of-concept that inhibiting the rRNA maturation process is as efficient as inhibiting rRNA synthesis to induce an antitumoral effect (Jouines et al., 2025). To determine whether rRNA maturation can be of therapeutic interest in neuroblastoma, we focused on neuroblastoma with unfavorable clinical outcome. In particular, we analysed the association between stage 4/4S neuroblastoma patient outcome and expression of two factors involved in rRNA maturation, fibrillarin (FBL) and WDR12.

We first analysed a test cohort using the Versteeg transcriptomic dataset (n=88) of the R2 database, using the median as a cut-off value. A significant association was observed between *FBL* mRNA levels and both neuroblastoma patients’ overall survival (OS) and relapse-free survival (RFS) (Figure 4A-B). Indeed, high *FBL* mRNA levels are associated with poor OS and RFS. Similar significant association was observed for WDR12 (Sup. Figure 1A). These data suggested that FBL and WDR12, two rRNA maturation factors, are prognosis marker in neuroblastoma.

**Figure 4.**
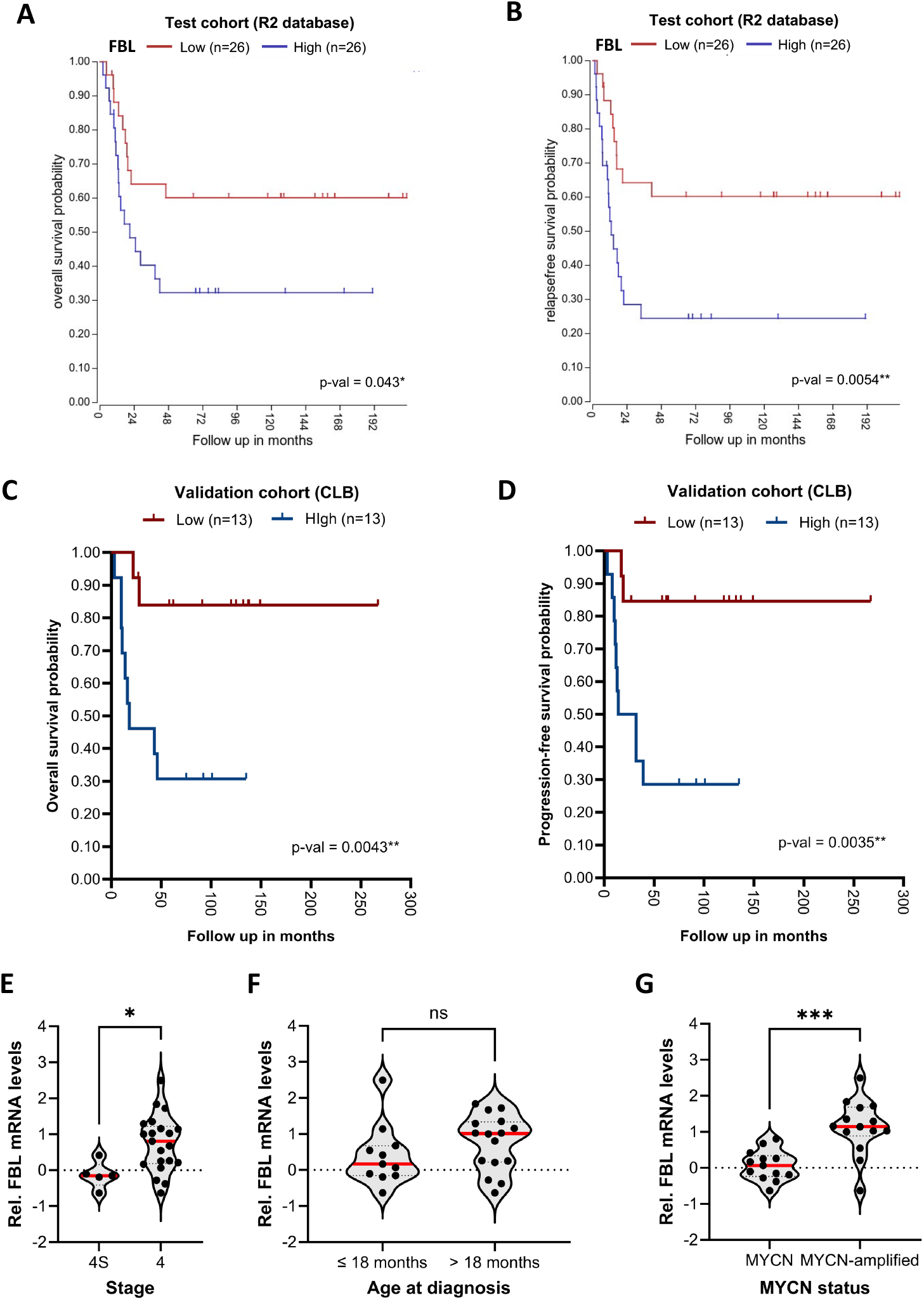
Prognostic value of *FBL* mRNA levels in neuroblastoma patients. **(A-B)** Test cohort. Association between *FBL* mRNA levels determined by microarray (R2 database) and overall survival (A) or relapse-free survival (B) was evaluated in 88 samples using Kaplan-Meier curves and log-rank Cox regression based on median as cut-off value. **(C-G)** Validation cohort. Association between *FBL* mRNA levels determined by medium-throughput RT-qPCR and overall survival (C) or progression-free survival (D) was evaluated in 26 samples using Kaplan-Meier curves and log-rank Cox regression based on median as cut-off value. FBL mRNA levels was compared in the validation cohort between 4S and 4 stage (E), early and late diagnosis (F), and MYC amplification status (G).

A validation cohort was then build using samples from the Centre Léon Bérard (CLB) (n=26). The validation cohort displays usual clinical characteristics of advanced neuroblastoma, including a poor outcome for neuroblastoma patients with stage 4, diagnosed after 18 months of age and with tumors displaying *MYCN*-amplification (Sup. Figure 2). Using medium-throughput RT-qPCR and median as a cut-off value, we observed that high *FBL* mRNA levels significantly associate with poor overall survival (OS) and progression-free survival (PFS) (Figure 4C-D). Similar significant association was observed for WDR12 (Sup. Figure 1A). Thus, test and validation cohorts demonstrated that high expression levels of factors involved in rRNA maturation are associated with poor outcome in neuroblastoma, supporting their potential as therapeutic target.

Interestingly, a significant correlation was observed between *FBL* and *WDR12* mRNA levels in the test cohort (Sup. Figure 1B). In addition, high expression of FBL and WDR12 are observed in neuroblastoma patients of the test cohort diagnosed lately and with tumors displaying *MYCN*-amplification (Sup. Figure 1B). Similar tendencies were observed in the validation cohort (Figure 4E-G and Sup. Figure 4C-E). In the validation cohort, *FBL* mRNA levels were significantly higher in stage 4 than stage 4S (Figure 4E), a non-significant tendency also observed for *WDR12* mRNA (Sup. Figure 1C). When regarding the age of neuroblastoma patients at diagnosis, similar mRNA levels of *FBL* and *WDR12* were observed before or after 18-month of age (Figure 4F and Sup. Figure 1D), despite the high predominance of stage 4 in older patients (Sup. Figure 2A). Interestingly, both *FBL* and *WDR12* mRNA levels were higher in *MYCN*-amplified tumors compared to non-amplified *MYCN* tumors, an observation consistent with the ribosome biogenesis promoting activity of MYC (Figure 4G and Sup. Figure 1E).

These findings suggest that factors involved in rRNA maturation are marker of poor prognosis. Compared to *WDR12, FBL* is over-expressed in tumors with the poorest outcome (i.e., stage 4, diagnosis > 18 months, *MYCN*-amplification), highlighting the clinical potential of FBL in neuroblastoma.

### FBL knock-down reduces cell proliferation in neuroblastoma

FBL overexpression being associated with poor outcomes in neuroblastoma, we evaluated the impact of inhibiting FBL on neuroblastoma cell proliferation. A stable IMR-32 cell line expressing an inducible shRNA targeting FBL was generated. FBL knock-down was validated by analysing FBL expression by Western blot in response to one week of induction with doxycycline (Figure 5A-B). In response to doxycycline treatment, FBL protein levels were reduced by 95% in the shFBL cell line, while no change in FBL protein levels was observed in the shNS control cell line.

**Figure 5.**
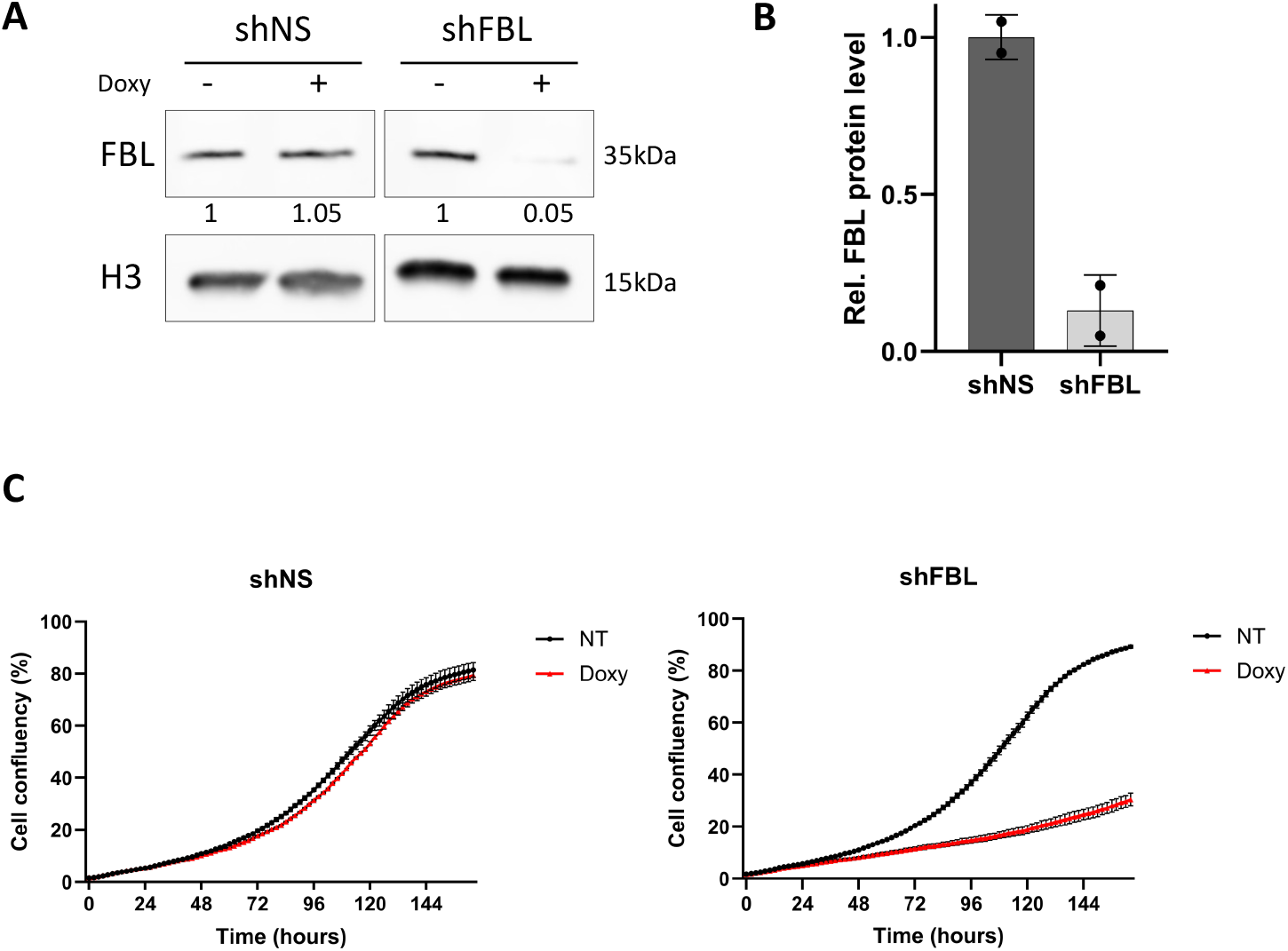
FBL knock-down reduces neuroblastoma cell proliferation. **(A-B)** Western blot of FBL detection in stable shRNA-derived IMR-32 cell lines, after induction with doxycycline (Doxy) at 1 µg/mL for one week. Cells express either a FBL-targeting shRNA (shFBL) or a non-targeting control shRNA (shNS). A representative gel is given in (A) and quantification of protein levels in (B). **(C)** Proliferation of shRNA expression IMR-32 was monitored over 10 days using real-time imaging in response to 1 µg/mL doxycycline induction. Data are mean +/- sd of 3 technical replicates.

Growth of shRNA-expressing IMR32 cell lines was monitored in real time for six days (Figure 5C). No significant difference in cell proliferation was observed between non-induced and doxycycline-induced shNS control cells (Figure 5C, left panel). In contrast, in shFBL cell line, FBL knock-down reduced cell proliferation as early as from 36 h post-induction. At the end-point (i.e., 144 h), shFBL expressing cells reached only 30% confluence compared to 90% for the shNS expressing cells (Figure 5C, right panel).

These data indicates that FBL inhibition reduces cell proliferation, supporting its potential as a therapeutic target in neuroblastoma.

## Discussion

Neuroblastoma, a prevalent paediatric cancer, presents significant clinical challenges, particularly in the case of high-grade tumors bearing *MYCN* amplification, where innovative therapeutic approaches are still needed. This study underscores the potential of targeting ribosome biogenesis pathways, including the rRNA maturation process, as a therapeutic approach in neuroblastoma with unfavorable clinical outcome.

Conflicting data has been published on the impact of MYCN on neuroblastoma cell sensitivity. While MYCN was found to sensitise cell lines to CX-5461 and CX-3543 [10], Several studies demonstrated that the RNA Pol I inhibitor, CX-5461, reduces neuroblastoma growth *in vitro* and *in vivo*, independently of *MYCN* amplification status [6], [14], [4]. Our data confirm the latter observations in a panel of neuroblastoma patient-derived cell lines established in our laboratory. The use of patient-derived cell lines adds clinical relevance to the efficacy of RNA Pol I inhibitors in neuroblastoma. In addition to CX-5461, we report that the BMH-21 RNA Pol I inhibitor also reduces neuroblastoma cell proliferation. Importantly, we and others reported that CX-5461 induce DNA damage and also acts as a topoisomerase 2 inhibitor [15], [13]. In contrast, BMH-21 did not induce any DNA damage, supporting that the overall vulnerability of neuroblastoma to BMH-21 relies solely on ribosome biogenesis inhibition. This study is also the first to demonstrate the efficacy of BMH-21 in neuroblastoma, which strengthens the potential of ribosome biogenesis inhibition as an anti-tumoral strategy. Both CX-5461 and BMH-21 induce a ribosomal stress response in neuroblastoma cell lines, as evidenced by the increased p21 expression and activation of the caspase 3/7 pathway. Our data supports that RNA Pol I inhibitors promote both cytostatic and cytotoxic effects, and further substantiates the role of ribosome biogenesis inhibition in curbing neuroblastoma proliferation [6], [10]. Interestingly, we recently reported that these two RNA Pol I inhibitors promote cell cycle arrest but not cell death in triple-negative breast cancer, suggesting that RNA Pol I inhibitors display distinct cellular response depending on the cancer type [13]. More importantly, these two RNA Pol I inhibitors show efficacy at nanomolar concentrations, indicating a high affinity of these compounds for their targets and reducing the risk of side effects in neuroblastoma patients.

Our data also demonstrate that other steps of ribosome biogenesis, precisely the rRNA maturation process, may be inhibited to exert an antitumoral effect on neuroblastoma cells. Among the factors involved in rRNA maturation, early foundational studies revealed the role of FBL in pre-rRNA maturation [16], [17], [18]. FBL is not only involved the rRNA-precursor processing but also in the catalysis of rRNA 2’O-ribose methylation, two maturation processes critical for proper ribosome assembly and function [19], [20], [18]. Using two independent cohorts of advanced neuroblastoma, we reveal that FBL is a prognostic marker in neuroblastoma, reporting a significant association between high FBL expression levels and poor overall and relapse-free survivals. The association of FBL with prognosis has been reported in other cancers such as breast cancer [21], [22]. Importantly, we demonstrated that prolonged FBL knockdown drastically reduces neuroblastoma cell proliferation, as we recently reported in triple-negative breast cancer [13]. *In vivo*, FBL inhibition induced a reduction in tumor frequency and volume, suppressing tumorigenesis [22], [13]. Overall, these findings support FBL’s potential as a therapeutic target in neuroblastoma.

Altogether, this study presents new opportunities for the management of paediatric neuroblastoma patients. Ribosome biogenesis inhibitors, whether through RNA Pol I or rRNA maturation factors inhibitors, emerges as a promising strategy to exploit the dependency of *MYCN*-amplified neuroblastoma cells on this process.

## Materials and methods

### Cellular models and treatments

The five patient-derived neuroblastoma cell lines, along with information on the *MYCN* amplification status, were developed and maintained by the V. Combaret group from paediatric neuroblastoma patients of Centre Léon Bérard (Table S1) [11]. The IMR32 cells and the five patient-derived cells were cultured in RPMI 1640 medium (LIFE TECHNOLOGIES) supplemented with 10% Fetal Bovine Serum (CLINISCIENCES) and 1% penicillin-streptomycin (LIFE TECHNOLOGIES). All cells were maintained at 37°C with 5% (v/v) CO_2_ in a humidified incubator.

Stable IMR-32 cell lines expressing inducible shRNAs were generated via lentiviral infection as described previously (Erales PNAS 2017). Briefly, lentiviral particles were produced using pTRIPZ-shRNA-NS and pTRIPZ-shRNA-FBLvectors (Open Biosystems, Horizon Discovery) (Table S2). Following lentiviral infection, cell populations were selected for 14 days with 1 µg/mL puromycin. shRNA expression was induced using 1 µg/mL doxycycline.

CX-5461 (orb251392, Biorbyt) was dissolved in a 25 mM NaH_2_PO_4_, - Na_2_HPO_4_ buffer as previously described [23]. BMH-21 (orb304073, Biorbyt) was dissolved in a 25 mM phosphate-citrate buffer as previously described [24].

### Neuroblastoma cohorts

The test cohort of advanced neuroblastoma corresponds to the Versteeg dataset publicly available on the R2 database (http://r2.amc.nl). It is composed of 88 samples and both transcriptomic and clinical data are available. The validation cohort of advanced neuroblastoma is composed of 26 samples (PGEB platform, Cancer Research Centre of Lyon/Centre Léon Bérard, Lyon, FRANCE). Informed written consent from parents, guardians, or patients was obtained for the use of biological samples and the use of data, and the study was approved by the Ethics Committee of Centre Léon Bérard in agreement with the Good Clinical Practice guidelines of the International Conference on Harmonization and the Declaration of Helsinki.

### Cellular based Assay

Cell proliferation was assessed using the CellTiter 96® AQueousOne Solution Cell Proliferation Assay kit (Promega, G3580) following the manufacturer’s instructions. Cell apoptosis was measured using the caspase-Glo® 3/7 reagent (Caspase-Glo® 3/7 Assay, Promega, G8091) following the manufacturer’s instructions. Absorbance values at 490 nm and luminescence were measured respectively, using a Tecan® Infinite 200 Pro plate reader. Cell proliferation was monitored using a real-time imaging system (Sartorius, Incucyte™ S3) on the Plateau d’Imagerie Cellulaire (PIC, Cancer research Center of Lyon). Cellular confluence was measured using the Incucyte software (Sartorius).

### RNA extraction and RTqPCR

Total RNA was extracted using the TRIzol-Chloroform method and quantified with the NanoDrop 2000 spectrophotometer (Thermo Scientific). Reverse transcription was performed on 200 ng of purified RNA using the PrimeScript RT kit (Takara, RR037B) with a mix of oligo dT primers and random 6-mer primers. Low-throughput real-time quantitative PCR (RT-qPCR) was carried out using the Light Cycler 480 SYBRGreen Master Mix (Roche, 4887352001) with the appropriate primer pairs (Table S3) on the LightCycler® 96 instrument. Medium-through put RTqPCR was performed as previously described using the BiomarHD system [21]. Each biological sample was analysed in triplicate. Relative fold-changes were calculated using the 2-ΔΔCT method [25].

### Western Blot

Proteins were extracted in Laemmli buffer and denatured for 10 minutes at 95°C. Protein quantification was performed using a TCA assay. Proteins were separated on a 4%-20% SDS-PAGE gradient gel and transferred onto a nitrocellulose membrane. Membranes were saturated in Tris Borate Saline buffer containing 0.05% Tween-20 and 5% skim milk and incubated with the primary antibodies for 1 hour at room temperature (H3, 1/5000, ab1791; Fibrillarin, 1/1000, ab166630). Secondary antibodies conjugated to horseradish peroxidase (Cell Signaling 1:5000) were used for chemiluminescence detection on a ChemiDoc MP (Bio-Rad) after incubation with Clarity ™ ECL (1705060, Bio-Rad), and signals were quantified using ImageLab software (Bio-Rad).

### Statistical analysis and graphical visualisation

Mean comparison was performed using the non-parametric Mann-Whitney test. Survival curves with associated log-rank tests were generated using the Kaplan-Meier method for overall survival (OS: from diagnosis to death), relapse-free survival (RFS: from diagnosis to either death or local recurrence) or progression-free survival (PFS: from diagnosis to either death or local/distant recurrence). Statistical analyses and graphical visualisations were performed using the R2 database (http://r2.amc.nl) and GraphPad Prism v7.0a (GraphPad Software, Inc).

## Declarations

### Ethics approval and consent to participate

Informed written consent from parents, guardians, or patients was obtained for the use of biological samples and the use of data, and study was approved by the Ethics Committee of Centre Léon Bérard in agreement with the Good Clinical Practice guidelines of the International Conference on Harmonization and the Declaration of Helsinki.

## Consent for publication

Not applicable

## Availability of data and material

The Versteeg dataset is publicly available on the R2 database (http://r2.amc.nl). The dataset generated from the Centre Léon Bérard biobank is available from the corresponding authors on reasonable request. All other data generated during the study are available in the manuscript and supplementary files, and are available from the corresponding authors upon request. Biological material is available upon reasonable request.

## Authors contributions

CJ, PLM, AG, M-AM, VS, DM, and IDI performed experiment and analysed data. VM and FC analysed data. IDI and VC contributed cellular models. VM, VC and FC conceived the study. FC, JJD and VM secured funding. CJ, VM and FC wrote and reviewed the manuscript.

## Acknowledgements

We thank the core facilities of the Cancer Research Center of Lyon and the associated staff not cited here for technical help: in particular Christophe Vanbelle and Mélina Gautier (PIC Cellular Imaging Platform of CRCL) and Séverine Tabone-Eglinger (PGEB biobank, Centre Léon Bérard). We thank Brigitte Manship for editing the manuscript.

