## Supplementary Figures for "Ribosome biogenesis is a therapeutic vulnerability in paediatric neuroblastoma"

Supplementary Figure 1

**A**

| WDR12 | Test cohort (R2 database) | Validation cohort (CLB) |
|---|---|---|
| Low (L) | n = 26 | n = 13 |
| High (H) | n = 26 | n = 13 |
| **Overall survival** | p-val < 0.000 1*** (L>H) | p-val = 0.042* (L>H) |
| **Relapse-free survival** | p-val < 0.000 1*** (L>H) | na |
| **Progression-free survival** | na | p-val = 0.033* (L>H) |

**B**

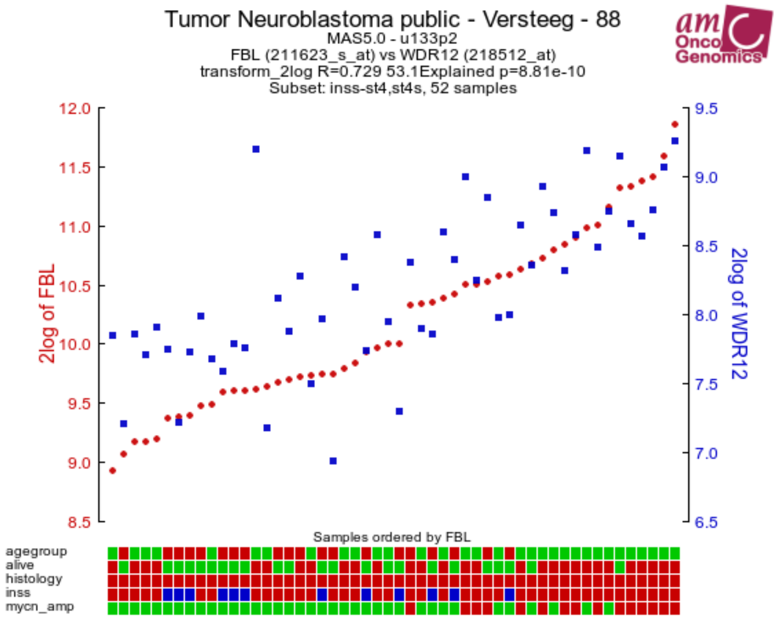


**C**

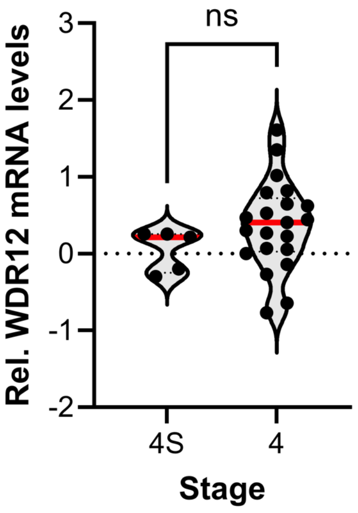


**D**

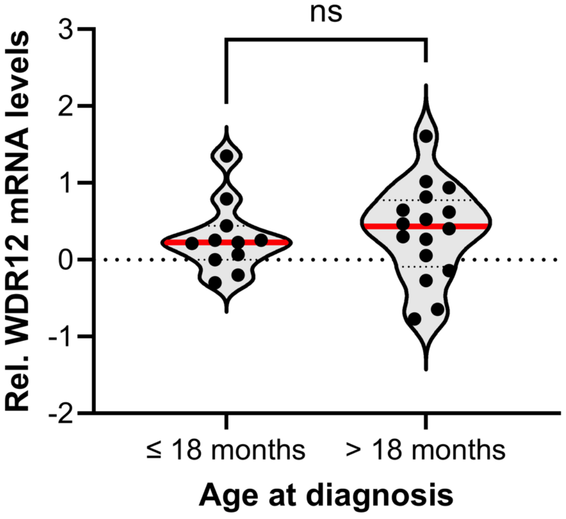


**E**

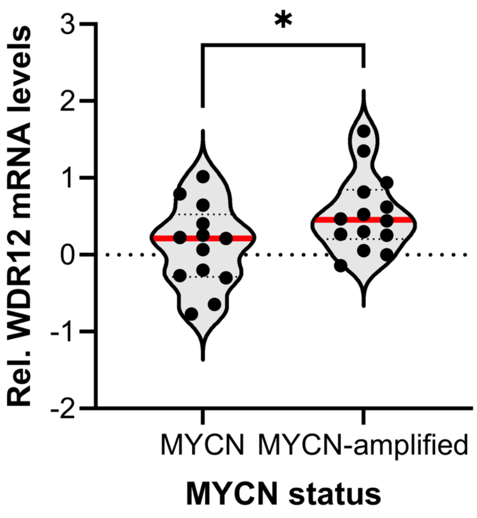

**Figure S1: Prognostic value of *WDR12* mRNA levels in neuroblastoma patients. (A)** Association between *WDR12* mRNA levels and overall, relapse-free or progression-free survivals was evaluated in both the test and validation cohorts. **(B)** Correlation of FBL and *WDR12* mRNA levels in the test cohort. **(C-E)** *WDR12* mRNA levels was compared between 4S and 4 stage (C), early and late diagnosis (D), and MYC amplification status (E) in the validation cohort.

### Supplementary Figure 2

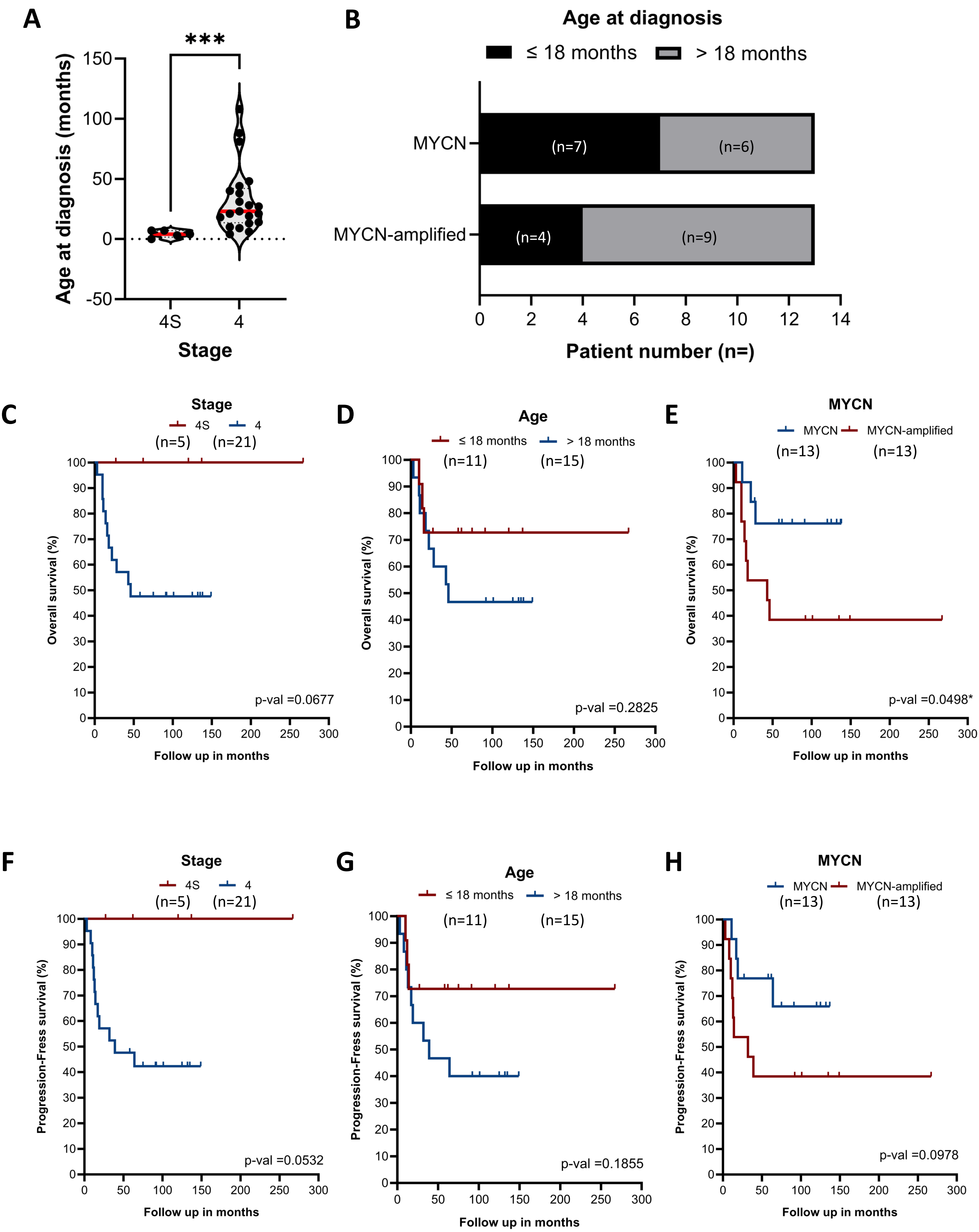

**Figure S2: Characteristics of the validation cohort. (A-B)** Increased of age at diagnosis in stage 4 compared to stage 4S (A) and in MYC-amplified tumors compared to normal ones (B). **(C-H)** Association between stage (C, F), age at diagnosis (D, G), MYC amplification status (E, H) and overall (C-E) or progression-free survivals (F-H).
