## Supplementary Tables for "Ribosome biogenesis is a therapeutic vulnerability in paediatric neuroblastoma"

*Table S1: MYC Amplifications in Neuroblastoma Cells*

| Name | ***MYCN*** **amplification** |
|---|---|
| CLB-LEC | yes |
| CLB-Ba | yes |
| CLB-BOULT | no |
| CLB-Ga | no |
| CLB-Sedp | no |
| IMR-32 | yes |

*Table S2: shRNA Sequences*

| Name | Sequence |
|---|---|
| shARN-FBL | AGACCATCCGGACCAACGA |
| shARN-NS | GCGATCTCGCTTGGGCGAGAGTAAGTA |

*Table S3: Sequences of Primers Used for PCR*

| ARN | Forward Primer | Reverse Primer |
|---|---|---|
| P21 | ACT CTC AGG GTC GAA AAC GG | CGG CGT TTG GAG TGG TAG AA |
| hTERT | TTC CGC AGA GAA AAG AGG GC | GTT GAG CAC GCT GAA CAG TG |
| FBL | CCT GGG GAA TCA GTT TAT GG | CCA GGC TCG GTA CTC AAT TTT |
| WDR12 | TAA AGG GGC AGA GGA ATG GAT | CAA CAT CCG TAT GTC CCA CAA |
| GAPDH | AGC CAC ATC GCT CAG ACA C | GCC CAA TAC GAC CAA ATC C |
| Actin | TCC ATC ACG ATG CCA GTG | CCA ACC GCG AGA AGA TGA |
| HPRT1 | TGA CAC TGG CAA AAC AAT GCA | GGT CCT TTT CAC CAG CAA GCT |
| PGK1 | AAG TGA AGC TCG GAA AGC TTC TAT | AGG GAA AAG ATG CTT CTG GG |
| PPIA | GTC AAC CCC ACC GTG TTC TT | CTG CTG TCT TTG GGA CCT TGT |
